# RIG-I immunotherapy overcomes radioresistance in p53-positive malignant melanoma

**DOI:** 10.1101/2021.10.16.464638

**Authors:** Silke Lambing, Stefan Holdenrieder, Patrick Müller, Yu Pan Tan, Christian Hagen, Stephan Garbe, Martin Schlee, Jasper G. van den Boorn, Eva Bartok, Gunther Hartmann, Marcel Renn

## Abstract

Radiation therapy induces cytotoxic DNA damage, which results in cell-cycle arrest and activation of cell-intrinsic death pathways, but its application has been limited by the radioresistance of tumors, such as in malignant melanoma. RIG-I is a cytosolic immune receptor expressed in all somatic cells, including tumor cells, with a key role in sensing viral RNA. RIG-I specific oligonucleotide ligands elicit a robust cell-intrinsic antiviral response and immunogenic cell death in tumor cells and are being tested in clinical trials. Nonetheless, their potential to overcome radioresistance has not yet been explored. Here, we demonstrate that activation of RIG-I enhances the extent and immunogenicity of irradiation-induced tumor cell death in human and murine melanoma cell lines *in vitro* and improved survival in the murine B16 melanoma model. Pathway analysis of transcriptomic data revealed a central role for p53 downstream of the combination treatment, which was corroborated using *p53*^*-/-*^ B16 cells. *In vivo*, the effect of irradiation on immune-cell activation and inhibition of tumor growth was absent in mice carrying *p53*^*-/-*^ B16 tumors, while the response to RIG-I stimulation in those mice was maintained. Our results identify p53 as pivotal for the synergistic antitumoral effect of RIG-I and irradiation, resulting in potent induction of immunogenic tumor-cell death. Thus, the administration of RIG-I ligands in combination with radiotherapy is a promising therapeutic approach to treating radioresistant tumors with a functional p53 pathway, such as malignant melanoma.

## Introduction

Radiation therapy is a mainstay of anti-tumor therapy, including lymphoma, breast, brain and head and neck cancers. It is currently used in the treatment schedule of 50% of all malignancies (1). Since radiotherapy has been demonstrated not only to restrict tumor-cell proliferation but also to induce tumor-specific CD8^+^ T-cells (2), it has also been studied in combination with tumor immunotherapies, such as checkpoint inhibitors, in pre-clinical and clinical trials (3). Mechanistically speaking, irradiated tumor cells have been reported to release pro-inflammatory cytokines, including CXCL16 and TNFα (4,5), the cGAS ligand cGAMP (6,7), and alarmins such as HMGB1 and ATP (8-10). There have also been reports that ionizing radiation (IR) can induce the expression of MHC class I proteins (11,12) and calreticulin (13,14) on the surface of irradiated, dying cells, which promotes recognition and internalization of the cells by phagocytes and subsequent T cell activation. However, many tumors, such as malignant melanoma, are primarily radioresistant or develop radioresistance upon repeated radiotherapy (15,16), limiting the utility of this approach.

Recent advances in immunotherapy have significantly prolonged survival for patients with many different tumor entities (17). While immune checkpoint inhibition is effective in a portion of patients, the presence of an anti-inflammatory (‘cold’) tumor microenvironment and lack of pre-existing tumor-antigen-specific T cells still poses strong limitations for checkpoint-inhibitor treatment in many patients (18). One promising approach for ‘converting’ the tumor microenvironment to make it amenable to immune-cell infiltration and mount an effective anti-tumor response is the targeted stimulation of innate immune receptors, including the cytosolic, antiviral receptor retinoic acid-inducible gene I (RIG-I). RIG-I is broadly expressed in nucleated cells, including tumor cells, and can be specifically activated by 5’-tri- or 5’-diphosphorylated, blunt-ended, double-stranded RNA (19-21). RIG-I activation leads to the induction of the antiviral mediator type I interferon (IFN) and pro-inflammatory cytokines (19-21). Moreover, it also directly induces tumor-cell death (22,23) that bears the classical ‘immunogenic’ hallmarks, such as HMGB1 release and calreticulin exposure on the cell surface (24-26). *In situ* RIG-I stimulation thus offers features of a cancer vaccine by simultaneously inducing the release of tumor antigens and creating a pro-immunogenic environment that facilitates the development of tumor-specific cytotoxic T cells (24,27).

We hypothesized that the combination of irradiation and specific RIG-I activation could change the tumor environment to ‘hot’ and thus enable an effective anti-tumor response. Herein, we investigated the combination of the synthetic RIG-I-specific ligand (3pRNA) with irradiation. We found that the combination of RIG-I activation and irradiation significantly increased immunogenic cell death in both human and murine melanoma cell lines and improved the uptake of dead tumor cells and the activation of dendritic cells. The analysis of transcriptomic data identified a critical role for the p53 pathway, which was confirmed by using *p53*^*-/-*^ B16 cells. While the anti-tumor effects of RIG-I monotherapy were independent of p53, the RIG-I-mediated increase in the susceptibility of tumor cells to irradiation was found to be p53-dependent. Cotreatment of 3pRNA with tumor-targeted irradiation resulted in increased activation of T-and NK cells in draining lymph nodes and prolonged the overall survival of tumor-bearing animals in an *in vivo* B16 melanoma model. Altogether, our study suggests that combined radio-RIG-I-immunotherapy has great clinical potential, especially in patients with radioresistant tumors exhibiting an intact p53 pathway, like most malignant melanomas have.

## Material & Methods

### Cell lines

Human A375 and SKmel28 melanoma cells, A549 lung adenocarcinoma cells, and murine B16.F10 melanoma cells were cultured in DMEM, and human melanoma cells MaMel19, MaMel54, and MaMel48 were cultured in RPMI 1640, both media were supplemented with 10% fetal-bovine-serum (FCS), 100 IU/ml penicillin, and 100 μg/ml streptomycin (all from Thermo Fisher Scientific) in a humidified incubator at 37°C and 5% CO_2_. A375 cells were kindly provided by Michael Hölzel (University Hospital Bonn, Germany), and MaMel19, MaMel54 and MaMel48 cells were kindly provided by Jennifer Landsberg (University Hospital Bonn, Germany) and Dirk Schadendorf (University Hospital Essen, Germany). B16 and Skmel28 cells were purchased from ATCC. The identity of the human cell lines was confirmed by short-tandem-repeat (STR) profiling (Eurofins). Cells were checked monthly for mycoplasma infection.

### Oligonucleotides, reagents, and chemicals

5’-triphosphorylated double-stranded RNA (3pRNA) was *in vitro* transcribed (IVT) from a DNA template by using the phage T7 polymerase from the Transcript Aid T7 high-yield Transcription Kit (Fermentas), as described previously (28). Inert AC_20_ control RNA (5’- CACAACAAACCAAACAACCA-3’) and polyA RNA were obtained from Biomers and Sigma Aldrich, respectively. Murine IFNα was purchased from BioLegend. The MDM2 inhibitor AMG232 was purchased from MedChemExpress.

### Oligonucleotide-transfection of tumor cells

Lipofectamine 2000 (Invitrogen) and OptiMem (Thermo Fisher Scientific) were used according to the manufacturer’s protocol to transfect control AC_20_ RNA or stimulatory 3pRNA at the indicated concentrations.

### Irradiation of tumor cells

Cells were irradiated with high-energy photons (150 keV) generated by a biological irradiator (RS-2000, Rad Source Technologies) 30 min after transfection of RNA or stimulation with IFNα.

### DC uptake of melanoma cells

Bone-marrow-derived dendritic cells (BMDCs) were generated from wildtype C57Bl/6 mice as described previously (29). B16 melanoma cells were stimulated as indicated in the figures. After 48 h, the melanoma cells were stained with eFluor780 fixable viability dye (eBioscience, 1:2000 in PBS) for 30 min on ice. Excess dye was washed away by the addition of 200μl DMEM supplemented with 10% FCS. Stained melanoma cells (25,000) were then cocultured in a 96-well plate with 100,000 BMDCs overnight. The next day, DCs were detached by adding 2 mM EDTA/PBS and analyzed by flow cytometry.

### Generation of polyclonal p53 knockout (KO) cell lines by using CRISPR/Cas9

The CRISPR target site for murine p53 (single guide (sg) RNA: 5’-CTGAGCCAGGAGACATTTTC-3’) was already cloned into a px330 plasmid (px330-U6-Chimeric_BB-CBh-hSpCas9, Addgene plasmid #42230) and for human p53 (sgRNA: 5’-GCATCTTATCCGAGTGGA-3’) was already cloned into a px459 plasmid (pSpCas9(BB)-2A-Puro (px459) V2.0 (Addgene plasmid #62988)) and kindly provided by Daniel Hinze from the lab of Michael Hölzel. B16 and A375 cells were seeded at a density of 5×10^4^ cells per well into a 12-well plate the day before transfection with 2 μg of the CRISPR/Cas9 plasmid using Lipofectamine 2000. After three days of incubation at 37°C, the transfected cells were seeded out again into 12-well plates at a density of 5×10^3^ cells per well. One day later, 10 μM of the MDM2 inhibitor AMG232 was added to the culture medium for five days to positively select for p53-deficient cells.

### Gene-expression analysis with microarray

B16.F10 cells were transfected with 50 ng/ml 3pRNA or AC_20_ control RNA and either irradiated with 2 Gy or not for 6 h. RNA was isolated with the RNeasy Mini Kit (Qiagen), according to the manufacturer’s instructions. The extracted RNA was further processed using a Clariom S Mouse Genechip (Thermo Fisher) at the LIFE & BRAIN Genomics Service Center Bonn.

### Western blot analysis

Total cellular protein was extracted as described previously (30). 30–50 μg of protein was mixed with an equal amount of 2x Laemmli buffer (200 mM Tris/HCl pH 6.8, 4% SDS, 20% glycerol, 200 mM DTT), denatured at 95°C for 5 min, separated by SDS gel electrophoresis (30 mA per gel, 1.5 h), and transferred onto a nitrocellulose membrane (GE Healthcare, 0.45 μm pore size of the membrane). Proteins were transferred to the membrane using 450 mA for 1.5 h. The membranes were blocked with 5% non-fat dry milk in TBST buffer (150 mM NaCl, 20 mM Tris, 0.1% Tween 20, pH 7.6) for 1 h at room temperature (RT) and incubated with the respective primary antibodies at 4°C overnight (anti-phospho-p53 (Ser15), anti-p53, anti-Puma, anti-p21 (all 1:1000, Cell Signaling)). HRP-coupled secondary antibodies, anti-rabbit and anti-mouse (Cell Signaling), were used 1:5000 or IRDye800-coupled anti-rabbit and anti-mouse (Li-cor Bioscience) antibodies were used 1:10,000 in 5% milk/TBST and incubated for 1 h at RT. Anti-actin-HRP antibody (Santa Cruz) diluted 1:5000 in 5% milk/ TBST or mouse/rabbit anti-ß-actin (Li-cor Bioscience) diluted 1:10,000 was used to detect actin as a loading control. Protein bands were detected by using chemiluminescence of an ECL western-blotting substrate (Thermo Scientific) or by near-infrared fluorescence with the Odyssey Fc (Li-cor Biosciences).

### Enzyme-linked immunosorbent assay (ELISA)

HMGB1 ELISA Kit from IBL International was used according to the manufacturer’s protocol.

### Flow cytometry

Cells of interest were harvested with trypsin and washed with PBS. For staining of surface proteins, fluorochrome-conjugated monoclonal antibodies were diluted 1:200 in FACS buffer (1x PBS containing 10% FCS, 2 mM EDTA and 0.05% sodium azide) and incubated with the cells 15–20 min on ice or RT. Antibodies used: APC-Cy7 or BV510 anti-CD4, PerCP-Cy5.5 or BV421 anti-CD8, PerCP anti-CD45, BV421 anti-CD11c, Alexa-Fluor-488 or BV510 anti-CD69, BV785 anti-CD86, BV785, BV510 anti-MHC-I (Hk2b), FITC anti-I-A/E (all BioLegend), FITC anti-CD11c, APC anti-MHC-I (Hk2b), PE or BV650 anti-NK1.1 (all eBioscience), BUV737 anti-CD4, BUV395 anti-CD8, BUV395 anti-CD11b, FITC anti-HLA ABC (all BD Bioscience), Alexa-488 anti-Calreticulin (Cell Signaling Technology, diluted 1:100).

For *in vivo* studies, the tissue was digested with 1 mg/ml collagenase D in PBS with 5% FCS for 20 min at 37°C and afterwards passed through a 70 μm cell strainer with PBS. Cells were stained with Zombie UV fixable viability stain (1:500 in PBS, BioLegend) for 20 min at RT, followed by blocking of Fc receptors (Anti-Mouse CD16/32 from eBioscience, 1:200 in FACS buffer) for 15 min on ice. Surface staining was performed as described above.

Intracellular staining of activated, cleaved caspase-3 was analyzed using a rabbit anti-cleaved caspase-3 monoclonal antibody (1:500, Cell Signaling Technology) followed by a second staining with FITC-anti-rabbit IgG (1:200, BioLegend). Both antibodies were diluted in FACS buffer supplemented with 0.5% saponin.

Fluorescence intensities for all of the flow-cytometry-based assays were measured with the LSRFortessa flow cytometer (BD Biosciences), or with the Attune NxT Flow Cytometer (Thermo Fisher).

### Quantification of apoptotic cell death

Cells were stained with Annexin V-Alexa 647 or Annexin V-Pacific Blue (both 1:30, BioLegend) in Annexin binding buffer (10 mM HEPES, pH 7.4; 140 mM NaCl; 2.5 mM CaCl_2_) and incubated at RT for 20 min in the dark. Cells were washed and resuspended in 200 μl 1x binding buffer. 5 μl of 7-amino-actinomycin D (7AAD, 50 μg/ml working solution in PBS, Thermo Fisher Scientific) was added to the stained cells 5–10 min before measurement.

### Multiplex cytokine assay

Cytokine levels were measured using human and mouse LEGENDplex bead-based multi-analyte flow assay kits, as described in the manufacturer’s manual. However, the assay was performed in a 384-well plate and the volumes adjusted accordingly.

### Cell-cycle-phase analysis

Analysis of cell-cycle phases was performed on cells that were fixed and permeabilized with 70% ethanol for one hour at RT. Cells were incubated for 30 min at RT with 10 μg/ml propidium iodide (PI) and 100 μg/ml RNase A in FACS buffer, and directly analyzed by flow cytometry. For simultaneous staining of activated caspase 3, the cultivation medium of cells seeded in 96-well plates was exchanged for 50 μl/well of staining solution, containing CellEvent Caspase3/7 Green ReadyProbes, according to the manufacturer’s protocol, and 100 μg/ml Hoechst 33342 (both Thermo Fisher Scientific) and incubated for 30–60 min at 37°C. The cells were then detached and analyzed by flow cytometry.

### *In vivo* studies with mice

8–12 week-old female C57BL/6 mice were obtained from Janvier and housed in individually ventilated cages in the House of Experimental Therapy (HET) at the University Hospital Bonn under SPF conditions. Sample size was calculated a priori with G*Power (31). All experiments were approved by the animal ethics committee. After at least 3 days of acclimatization, mice were injected subcutaneously into the right flank of the back with 1×10^5^ B16.F10 cells in 100 μl sterile PBS. Mice with no tumor at the start of the experiment and mice with a tumor over 4 mm in diameter at the start of a survival or tumor-size experiment were excluded. When the tumors reached a diameter of 3–4 mm, the tumors were injected with 20 μg 3pRNA or control RNA complexed with *in vivo*-jetPEI (Polyplus) according to the manufacturer’s protocol and afterwards locally irradiated with a single dose of 2 Gy. For local irradiation, the mice were narcotized and positioned in the treatment beam. The tumors were stereotactically irradiated with adapted field size in a range between 1–2 cm using a linear accelerator with a 6 MeV beam (TrueBeam STx, Varian and Mevatron MD, Siemens). The mice were surrounded by water-equivalent RW3 sheets (PTW, Freiburg) and placed in the depth-plane Dmax (15 mm) of the 6 MeV-Beam. The tumor size was measured daily with a caliper and the volume calculated with the formula V = (W^(2)^ × L)/2. For the survival studies, mice having tumors with a diameter exceeding 10 mm had to be euthanized for ethical reasons.

### Statistical analysis

If not indicated otherwise, data are presented as the mean +/-SEM of at least three experiments. Normal distribution of the data was tested with the Shapiro-Wilk test. A statistical analysis of the difference between groups using t-test, one or two-way ANOVA, or Kruskal-Wallis test as appropriate and stated in the figure legends, was calculated with GraphPad Prism 9. * (P < 0.05), ** (P < 0.01), *** (P < 0.001), **** (P < 0.0001), ns: not significant.

## Declarations

### Ethics approval and consent to participate

All animal experiments were approved by the local authorities (LANUV NRW).

## Results

### Combined radio-RIG-I-immunotherapy induces immunogenic tumor cell death and tumor cell uptake by dendritic cells as well as activation in vitro

To investigate whether RIG-I activation combined with irradiation has a synergistic effect on the induction of immunogenic cell death *in vitro*, we stimulated the murine B16 and human A375 melanoma cell lines with the RIG-I ligand 3pRNA (28) followed by 2 Gy of ionizing radiation (IR). RIG-I synergistically increased irradiation-induced cell death, as measured by Annexin V positive and Annexin V/7AAD double-positive cells; notably, this effect could not be recapitulated by the addition of recombinant IFN-α to 2 Gy irradiated cells (Fig. 1 A-B, suppl. Fig. 1 A-B). Increased induction of cell death was confirmed by staining intracellular levels of cleaved caspase 3 (suppl. Fig. 1 C, D). Moreover, RIG-I activation and irradiation significantly lowered the EC_50_ for the induction of cell death as quantified by Annexin V/7AAD staining, from 987 ng/ml for 3pRNA alone to 293 ng/ml of 3pRNA in combination with 2 Gy radiation in murine B16 cells and from 1754 ng/ml to 333 ng/ml in human A375 melanoma cells (Fig. 1 C, D, suppl. Fig. 1 D, E). Since combined RIG-I stimulation had the greatest effect on cell death at 2 Gy (suppl. Fig. 1 F), this irradiation dose was selected for all subsequent experiments to analyze the effect of combination therapy. In addition to A375, we tested several other human melanoma cell lines (MaMel19, MaMel54, and MaMel48) and A549 lung adenocarcinoma cells, which also all showed increased cell death when RIG-I stimulation was combined with irradiation (Fig. 1 E, F).

**Figure 1:**
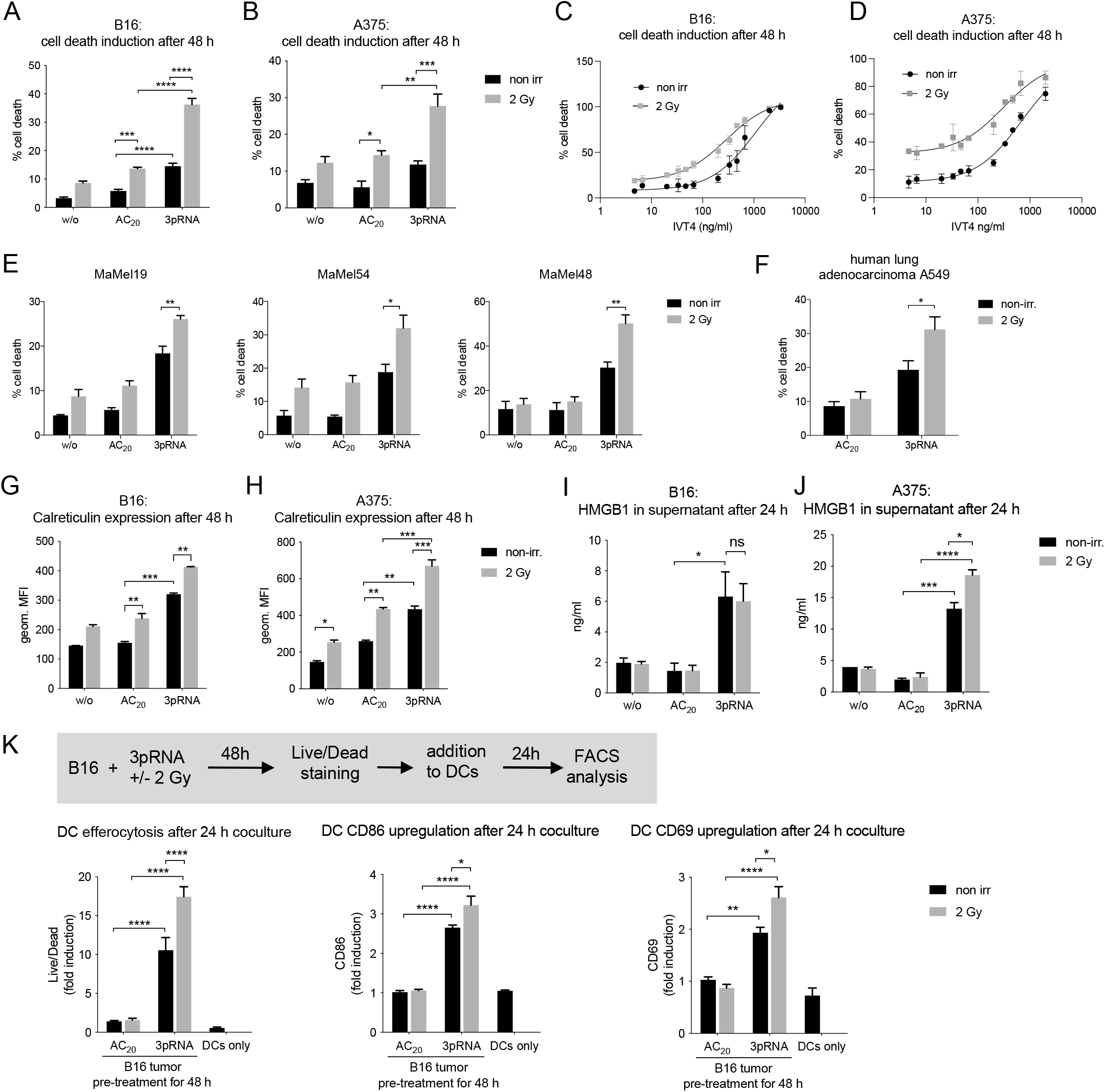
Irradiation enhances 3pRNA-induced immunogenic cell death in melanoma cells, as well as uptake by and co-stimulation of dendritic cells. (A-D) Murine B16 and human A375 melanoma cells were transfected with 50ng/mL 3pRNA or AC_20_ control RNA followed by 2 Gy irradiation. 48 h later, apoptosis was measured in B16 (A,C) and A375 (B,D) cells by using Annexin V/7AAD detection by flow cytometry. The dose of 3pRNA ligand was titrated in B16 (C) and A375 (D) cells to determine the EC_50_ values with and without 2 Gy. (E) Different human melanoma cell lines were transfected with 50 ng/ml (MaMel19) or 200 ng/ml (MaMel54, MaMel48) 3pRNA, and (F) human lung carcinoma cell line A549 was transfected with 50 ng/ml 3pRNA and irradiated (2 Gy) where indicated. Induction of cell death was quantified 48 h later using Annexin V/7AAD staining and flow cytometry. (G-J) Melanoma cells were transfected with 50 ng/ml 3pRNA and irradiated with 2 Gy. (G, H) expression of calreticulin on the cell surface after 48 h was measured by flow cytometry and (I, J) HMGB1 concentration in the supernatant after 24h was measured by ELISA. (K) B16 cells were treated with 200 ng/ml 3pRNA and 2 Gy for 48 h, stained with fixable viability stain, and cocultured with bone-marrow-derived DCs from wildtype BL/6 mice for 24 h. DC tumor-cell uptake and activation was measured by flow cytometry. % cell death was plotted as the sum of Annexin V^+^, Annexin V/7AAD^+^, and 7AAD^+^ populations divided by the total number of cells. A, B, E, F, K: data are shown as the mean and SEM of n=3 and I, J: n=2 independent experiments. C, D, G, H: Representative results shown with the mean and SD of n=3 independent experiments with similar results. * p<0,05; **p<0,01; ***p<0,001; ****p<0.0001; two-way ANOVA. w/o: untreated, AC_20_: control RNA, 3pRNA: 5’-triphosphate RNA, non-irr: non-irradiated.

Calreticulin exposure on the outer leaflet of the cell membrane induces the efferocytosis of dead or dying cells by antigen presenting cells (APCs) and is a hallmark of immunogenic cell death (32). In agreement with the Annexin V data, calreticulin exposure was also found to be significantly increased when irradiation and RIG-I activation were combined in murine B16 melanoma cells and human A375 cells (Fig. 1 G, H). Surface expression of calreticulin was highest in Annexin V/7AAD double-positive cells, which are known to be in the late stage of programmed cell death (suppl. Fig. 1 G). Interestingly, the expression of MHC-I on murine B16 cells and human A375 melanoma cells was also strongly induced by the combination treatment, most prominently on Annexin V/7AAD-negative cells (suppl. Fig. 1 G, H, I). Furthermore, the release of the nuclear protein HMGB1, which serves as a danger-associated molecular pattern (DAMP) and is characteristic of immunogenic cell death, was induced by RIG-I stimulation in both cell lines and further increased by 2 Gy irradiation in human A375 cells (Fig 1 I, J). RIG-I stimulation, but not 2 Gy irradiation, induced the release of type I interferon in murine B16 melanoma cells and types I and III interferon in human A375 cells. In murine B16 cells, combination treatment slightly enhanced the secretion of IL6 and TNFα but did not lead to an increase in the release of interferons or the interferon-stimulated chemokine CXCL10 (suppl. Fig. 2 A), whereas in human A375 cells, IL6, GMCSF, IL29 (interferon lambda-1) and CXCL10, but not IFN-β, was enhanced by irradiation when added to RIG-I stimulation (suppl. Fig. 2 B).

**Figure 2:**
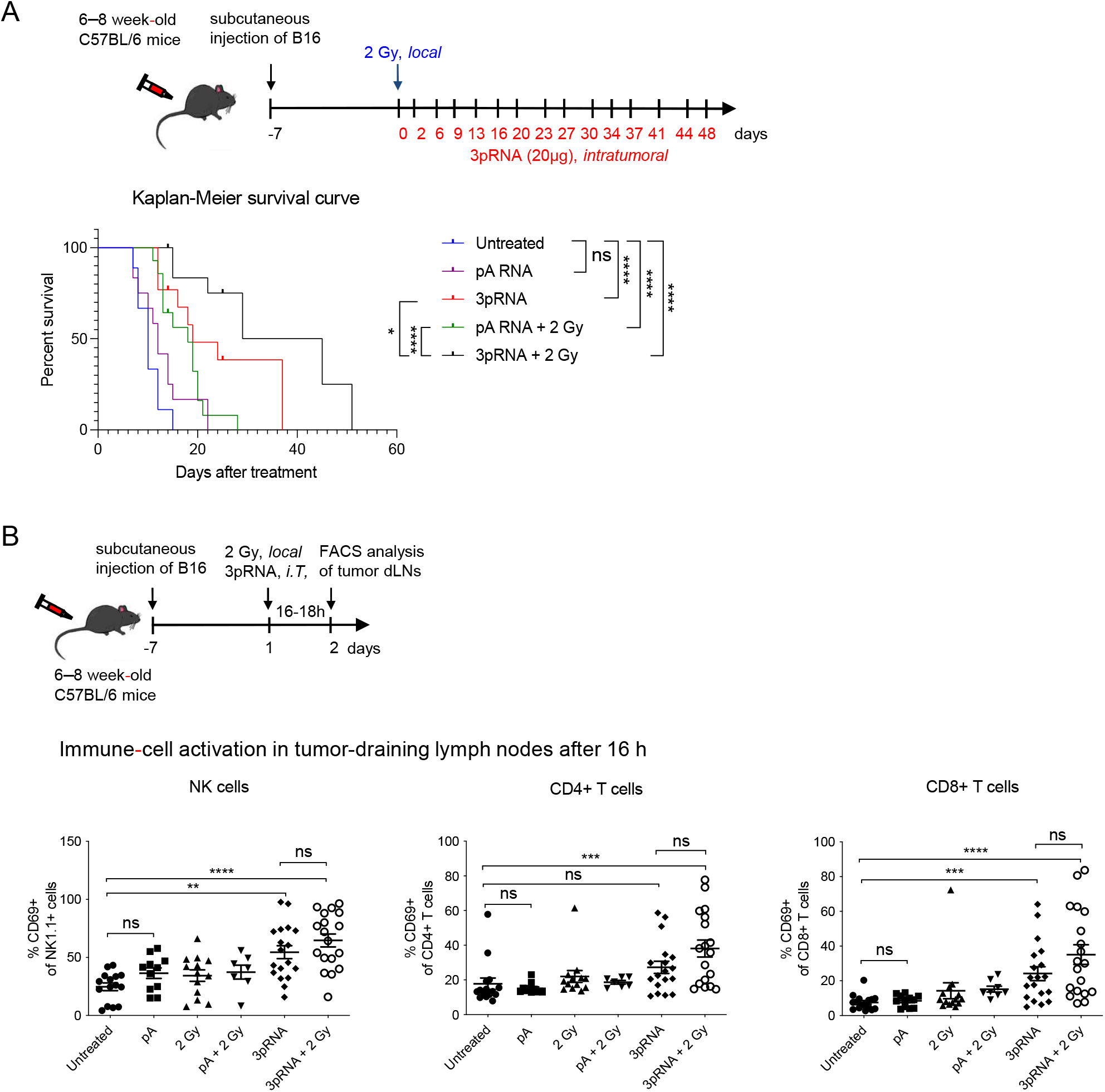
Concurrent irradiation and RIG-I immunotherapy prolongs the survival of melanoma-bearing mice. (A) B16 melanoma cells, subcutaneously transplanted into C57/BL6 mice, were locally irradiated with 2 Gy, injected with 20 μg 3pRNA, 20 μg control RNA (pA) or a combination of both, as indicated, and tumor size was measured regularly over 49 days. Mice with tumors larger than 10 mm in diameter were euthanized for ethical reasons. Survival rate is shown as a Kaplan−Meier curve. Summary of 3 independent experiments with 3−5 mice per group and experiment. Mantel−Cox test. (B) Subcutaneously transplanted B16 cells were treated as indicated and approximately 16 h later immune cells from the tumor-draining lymph nodes were analyzed for the activation marker CD69. Mean ± SEM of n = 3 with 3−5 mice per group and experiment. Kruskal−Wallis test. ns, not significant; **p<0,01; ***p<0,001; ****p<0.0001; pA: control RNA, 3pRNA: 5’-triphosphate RNA

To test whether the combination treatment had an impact on tumor-cell uptake by professional antigen-presenting cells and their activation, B16 melanoma cells were treated as before with 3pRNA and irradiation, but then stained with the eFluor780 fixable live/dead dye and co-incubated with bone marrow-derived dendritic cells (BMDCs). BMDCs “fed” with B16 cells after combination treatment demonstrated higher levels of eFluor780 dye uptake than after irradiation or RIG-I activation alone (Fig. 1 K). Combination treatment also significantly enhanced the expression of the costimulatory molecule CD86 and the immune-cell activation marker CD69 (Fig. 1 K).

### *Radio-RIG-I-immunotherapy prolongs the survival of B16 melanoma-bearing mice* in vivo

Next, we studied the combination of irradiation and RIG-I activation *in vivo*. C57BL/6 mice with a palpable subcutaneous B16 melanoma were treated with 2 Gy precision irradiation of the tumor area and intratumoral injection of 20 μg 3pRNA or 20 μg of non-stimulatory polyA control RNA twice a week. Compared to untreated tumors, both 3pRNA treatment alone and treatment with irradiation and inert RNA prolonged the survival of the mice. However, the combination of irradiation and RIG-I activation resulted in the longest overall survival (Fig. 2 A). In tumor-draining lymph nodes, analyzed at 16 hours after treatment, NK cells and CD8^+^ T cells showed increased expression of the activation marker CD69 upon RIG-I activation, with the highest expression when RIG-I activation and irradiation were combined. In CD4^+^ T cells, only the combination treatment of RIG-I activation and irradiation induced significant upregulation of CD69 (Fig. 2 B).

### Transcriptomic analysis of melanoma cells after combination therapy reveals activation of the p53 signaling pathway

To explore the potential molecular mechanisms of the combination therapy, we performed whole-genome transcriptional analysis with an Affymetrix gene chip on B16.F10 cells six hours after treatment with 3pRNA and irradiation. Following RIG-I stimulation, we observed a strong change in gene-expression patterns and a robust induction of interferon-stimulated genes (ISGs), whereas irradiation alone primarily induced genes associated with the DNA damage response (Fig 3 A). As expected, a pathway analysis of differentially expressed genes showed that RIG-I stimulation was associated with pathways involved in innate immunity, while irradiation induced genes of the p53 pathway. The p53 pathway was also among the most significantly upregulated pathways in the combination group (Fig. 3 B) and the only differentially regulated pathway between RIG-I activation alone and its combination with irradiation (Fig. 3 C, D). Given the central role of p53 signaling in DNA damage and cell-cycle control, we reasoned that this pathway may also be involved in the synergistic antitumoral effects observed for the combination treatment.

**Figure 3:**
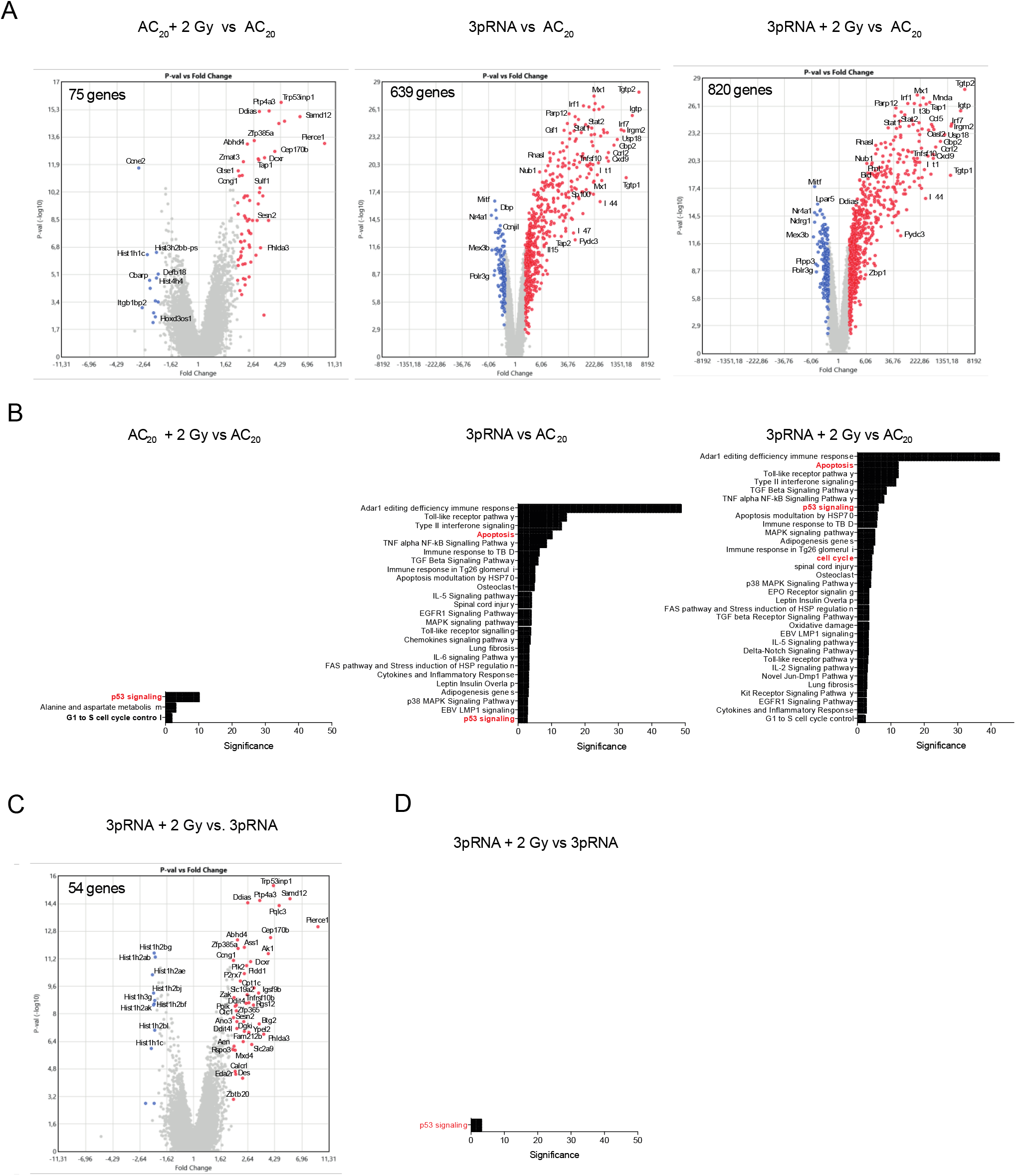
Whole-genome transcriptional analysis of B16 cells treated with the combined RIG-I radio-immunotherapy reveals activation of p53 signaling. Gene-expression analysis (Affymetrix GeneChip) of B16 total RNA 6 h after stimulation with 50 ng/ml 3pRNA or AC_20_ control and 2 Gy irradiation alone or in combination. (A) Volcano plots of single treatments and combined treatment in comparison to the control-transfected B16 cells or (C) combined treatment vs. 3pRNA-transfected cells. Colored data points show up- (red) or down-(blue) regulation of at least a 2-fold change. FDR corrected p-value < 0.05 (B,D) Pathway analysis (Wikipath) of genes found in (A) and (C) using the TAC software of Thermo Fisher ordered by significance. AC_20_: control RNA, 3pRNA: 5’-triphosphate RNA.

### Combined irradiation and RIG-I activation synergistically induces p53 signaling and prolongs cell-cycle arrest

We then examined the effect of RIG-I activation, irradiation, and combination treatment on p53 phosphorylation and signaling. As expected, irradiation induced p53 phosphorylation six hours after treatment, which then declined after 24 hours (Fig. 4 A). In contrast, RIG-I activation alone only led to weak p53 phosphorylation and only after 24 hours. However, combination treatment with 3pRNA and irradiation caused B16 cells to retain strong p53 phosphorylation even 24 hours after treatment (Fig. 4 A). Notably, total p53 protein levels at 24 hours were only elevated in 3pRNA-transfected B16 cells (both with and without irradiation). Moreover, these effects were not seen when irradiation was combined with control RNA or IFNα. We then analyzed the expression of two target proteins induced by p53, proapoptotic PUMA and the cell-cycle inhibitor p21 24h after treatment (Fig. 4 B). PUMA as well as p21 were induced by RIG-I activation and irradiation, with the strongest signal in the combination group, showing that the effects seen on p53 stability and phosphorylation (Fig. 4 A) translate into increased downstream effector molecule expression (Fig. 4 B).

**Figure 4:**
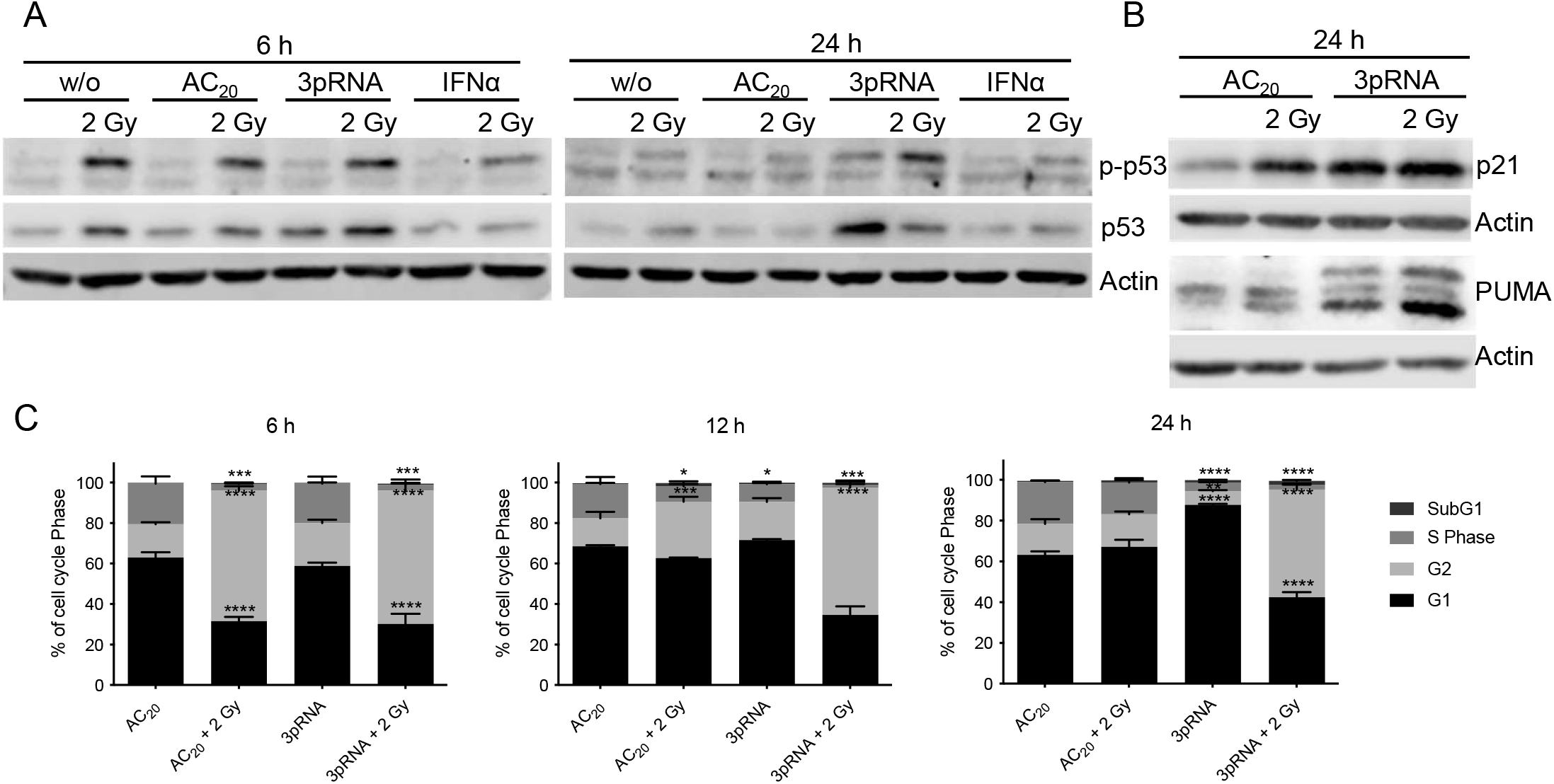
Combined RIG-I radio-immunotherapy induces p53 pathway activation and prolongs cell-cycle arrest. Western-blot analysis of (A) phospho- and total-p53 protein and (B) p21and PUMA expression after irradiation with 2 Gy, transfection of 50 ng/ml 3pRNA, or the combination of both in B16 cells at the indicated time points. Actin served as a protein-loading control. (C) Flow-cytometric cell-cycle analysis of B16 cells stained with propidium iodide and treated with 50 ng/ml 3pRNA and/or 2 Gy after the indicated time points. Mean and SEM of n=2. ns, not significant; * p<0,05; **p<0,01; ***p<0,001; ****p<0.0001; two-way ANOVA. AC_20_: control RNA, 3pRNA: 5’-triphosphate RNA.

To monitor effects on cell-cycle progression, we stained B16 melanoma cells with propidium iodide at 6, 12, and 24 hours after 2 Gy irradiation and RIG-I stimulation. Irradiation induced a G2/M cell-cycle arrest after 6 hours, which was already less pronounced after 12 hours and had been completely resolved 24 hours post-irradiation (Fig. 4 C). RIG-I stimulation alone, on the other hand, led to a G1/S arrest, which took 24 hours to develop, in line with its slower induction of p53 phosphorylation when compared to irradiation (Fig. 4 A). Like irradiation alone, the combination of irradiation and RIG-I stimulation led to a G2/M arrest after six hours. However, this arrest was maintained even after 24 hours (Fig. 4 C), which was consistent with the time course observed for p53 phosphorylation (Fig. 4 A).

### Synergistic effect of irradiation and RIG-I activation is p53-dependent, while the RIG-I effect alone is p53-independent

To test the functional relevance of p53 in combination therapy, we generated polyclonal p53-knockout (KO) cells using CRISPR/Cas9 genome editing. Polyclonal p53^-/-^ B16 and p53^-/-^ A375 melanoma cells showed no basal p53 expression and, as expected, following irradiation did not upregulate p53 protein at two hours or the p53 target protein p21 at 24 hours (Supp. Fig. 3 A-D). While the proportion of cell death induced by 3pRNA treatment alone was similar between wildtype and knockout cells, the additional increase upon irradiation observed in the WT cells was largely reduced in the p53^-/-^ cells (Fig. 5 A, B). In contrast to A375 cells carrying wildtype p53 (Supp. Fig. 3 E, left panel), there was no contribution of irradiation to cell-death induction in human p53-deficient SK-Mel28 melanoma cells, which carry an endogenous inactivating p53 mutation (33). Nevertheless, despite the lack of functional p53, RIG-I stimulation still strongly induced cell death in SK-Mel28 (Supp. Fig. 3 E, right panel). Similar to what was observed for cell death, the G1/S arrest induced by RIG-I stimulation alone was still present in the p53^-/-^ B16 melanoma cells after 24 hours. Furthermore, in the p53^-/-^ cells, irradiation still induced G2/M arrest after six hours. However, combination treatment did not induce the prolonged G2/M arrest for 24 and 48 hours that was observed in WT cells (Fig. 5 C).

**Figure 5:**
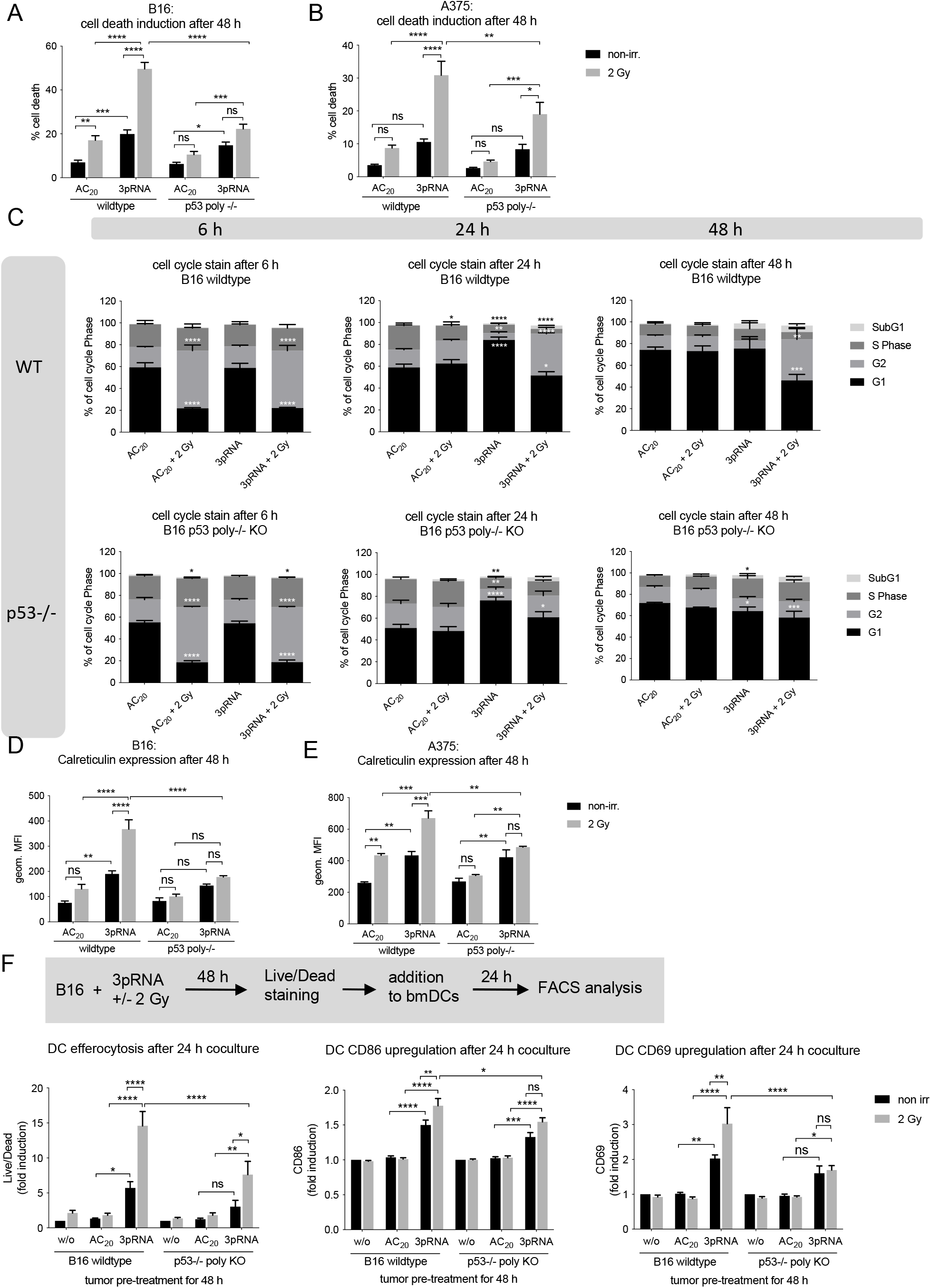
Knocking out p53 reduces the response of melanoma cells to combination treatment. (A−E) B16 or A375 wildtype and p53 polyclonal KO cells were transfected with 50 ng/ml 3pRNA, AC_20_ control RNA, or these in combination with 2 Gy irradiation. (A,B) Induction of cell death was quantitated via Annexin V/7AAD staining and analyzed by flow cytometry in B16 (A) and A375 (B) cells. (C) Flow-cytometric cell-cycle analysis with Hoechst 33342 at the indicated time points in B16 cells. (D,E) Surface calreticulin expression of B16 (D) and A375 (E) cells was monitored 48 h after treatment by flow cytometry. (F) B16 wildtype and p53^-/-^ cells were transfected with 200 ng/ml 3pRNA and irradiated as indicated 2 Gy. 48 h later cells were stained by Live-Dead eFluor780 stain and cocultured with bone-marrow derived DCs overnight. Activated DCs were analyzed by flow cytometry the next day. All data are shown as the mean and SEM of n=10 (A), n=5 (D), or n=3 (B, C, E, F). * p<0,05; **p<0,01; ***p<0,001; ****p<0.0001; two-way ANOVA. ns: not significant, w/o: untreated, AC_20_: control RNA, 3pRNA: 5’-triphosphate RNA, non-irr: non-irradiated

Analysis of caspase 3-positive cells in the individual phases of the cell cycle (G1, S, G2) showed that nearly all of the wildtype cells in the G2 phase after combination treatment undergo cell death after 48 hours (Supp. Fig. 4). Together with the lower proportions from other phases, there is a total of 60% caspase-3 positive cells for combination treatment, confirming the results of the Annexin V/7AAD staining (Fig 5 C) and underscoring the close link between cell-cycle arrest and cell death. Accordingly, in the absence of p53, additional irradiation had no significant effect when compared to RIG-I activation alone (Supp. Fig. 4). Moreover, for all phases of the cell cycle, the proportion of caspase-3 positive cells was substantially and significantly decreased for p53^-/-^ cells as compared to p53-wildtype cells, supporting the conclusion that the additional effect induced by irradiation requires functional p53 (Supp. Fig. 4).

Calreticulin expression on the cell surface of p53^-/-^ murine B16 or p53^-/-^ human A375 melanoma cells was also not further enhanced by combining RIG-I stimulation with irradiation (Fig. 5 D, E). Corresponding to the level of cell-surface calreticulin, the effect of irradiation on the uptake of p53^-/-^ B16 melanoma cells by DCs was markedly reduced in comparison to wildtype cells. Furthermore, no irradiation-dependent increase in the expression of the activation markers CD86 und CD69 on dendritic cells could be detected when the phagocytosed tumor cells lacked p53 (Fig. 5 F). This shows that the irradiation-dependent effects, including cell death, immunogenicity, subsequent uptake of dying cells by DCs, and activation of DCs, are primarily dependent on the expression of p53 in melanoma cells, whereas the effect of RIG-I treatment alone is not affected by the absence of p53.

### Synergistic anti-tumor activity of irradiation and RIG-I in vivo depends on functional p53 in melanoma

In the *in vivo* B16 melanoma model, both T cell activation and NK cell activation in the draining lymph nodes were enhanced by 3pRNA injection compared to untreated mice, as measured by upregulation of CD69 on CD8^+^ T cells, CD4^+^ T cells, and NK1.1^+^ NK cells (Fig. 6A). The addition of irradiation at the tumor area further enhanced the expression of activation markers on T cells in the draining lymph nodes. This additional irradiation-dependent stimulatory effect was lost in mice in which tumors were induced by injecting p53^-/-^ B16 melanoma cells (Fig. 6 A). These findings recapitulate the results obtained for immunogenic cell death and dendritic cell activation *in vitro* (Fig. 5). Consistent with activation of T cells and NK cells in draining lymph nodes, tumor growth was significantly reduced by RIG-I stimulation in wildtype and p53^-/-^ melanomas, and further reduced by additional local tumor irradiation. This additive effect was diminished and no longer statistically significant in p53^-/-^ tumors (Fig. 6 B), further supporting the notion that the synergistic effect of combination treatment *in vivo* is dependent on cell-intrinsic p53 expression in tumor cells. Nonetheless, the efficacy of RIG-I monotherapy is independent of the p53 status of the melanoma cells.

**Figure 6:**
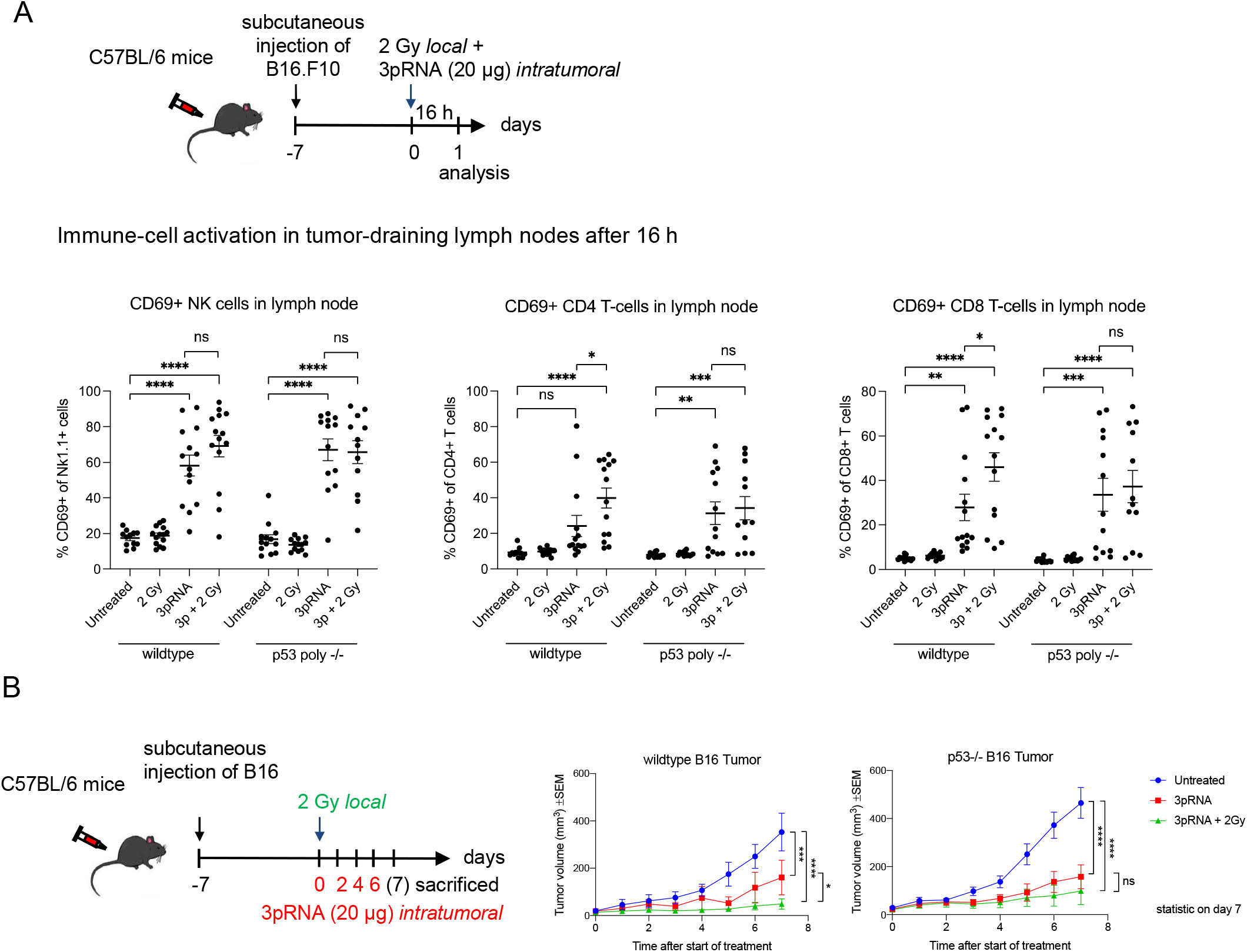
Synergistic anti-tumor activity of irradiation and RIG-I in vivo depends on functional p53 in melanoma. (A) B16 melanoma wildtype or p53 polyclonal knockout cells were subcutaneously transplanted into C57/BL6 mice and then locally irradiated with 2 Gy, injected with 20 μg 3pRNA, or a combination of both. 16 h later the mice were sacrificed. Tumor-draining lymph nodes were analyzed by flow cytometry for CD69 surface expression of activated CD8^+^ T cells, CD4^+^ T cells, and NK1.1^+^ NK cells. Mean and SEM of n=3 with 3–5 mice per group and experiment. (B) Mice were treated as indicated over 7 days and the tumor size was measured daily. Mean and SEM of n = 3 with 3–5 mice per group and experiment. ns, not significant; * p<0,05; **p<0,01; ***p<0,001; ****p<0.0001; two-way ANOVA. 3pRNA: 5’-triphosphate RNA

## Discussion

Through our studies, we found that a combination of RIG-I treatment with radiotherapy is a highly promising combinatorial treatment for tumors with an intact p53 pathway, such as most malignant melanomas (34). Localized irradiation of the tumor in a melanoma model *in vivo* substantially improved the therapeutic efficacy of intratumoral RIG-I ligand injections. This enhanced anti-tumor effect was accompanied by increased activation of CD4^+^ and CD8^+^ T cells in tumor-draining lymph nodes. *In vitro*, low-dose ionizing irradiation of tumor cells synergistically enhanced RIG-I-mediated induction of immunogenic tumor cell death, as characterized by increased cell-surface expression of calreticulin and the release of HMGB1 and inflammatory chemokines and cytokines. The uptake of this immunogenic material by dendritic cells caused them increasingly to be activated. Molecularly, the synergistic effect of irradiation and RIG-I could be ascribed to distinct effects on the p53 pathway, resulting in a prolonged cell-cycle arrest of tumor cells in the G2/M phase, which only occurred if RIG-I and irradiation were combined and which led to subsequent immunogenic cell death. Notably, the p53 pathway was required for synergistic activity *in vitro* and *in vivo* but not for the anti-tumor activity of intratumoral RIG-I ligand treatment as a monotherapy.

In approximately 50% of all human tumors, p53 is either mutated or functionally inactive (35) or MDM2 is overexpressed and downregulates p53 expression (36). Therefore, our data demonstrating that RIG-I therapy is independent of p53 are encouraging for RIG-I-mediated immunotherapy in general. Furthermore, based on our results, the combination of RIG-I with radiotherapy should be limited to treating tumors with an intact p53 pathway. In melanoma, the frequency of p53 mutations is only 10 to 19% (34), suggesting that the combination therapy is well suited to target malignant melanoma.

It is interesting to note that there is evidence from previous studies that p53 signaling is important to antiviral defense and interferon signaling (37,38). Moreover, it has been reported that treatment with IFN-β concurrent to irradiation or chemotherapy in mouse embryonic fibroblasts and in human hepatic cancer cells sensitized the cells for a higher induction of apoptosis (38). However, in our study, recombinant type I IFN was not a sufficient substitute for RIG-I stimulation, since it did not trigger enhanced and prolonged p53 phosphorylation or the induction of immunogenic cell death by radiotherapy.

To date, two studies have examined the combination of the non-specific antiviral receptor agonist poly(I:C) with radiation (39,40). However, it should be noted that poly(I:C) activates multiple dsRNA receptors, including PKR, OAS1, ZBP1, TLR3, MDA5, and RIG-I (41) rendering this rather non-specific immunotherapeutic approach more prone to interindividual variability and immunotoxic side effects. In one study, the combination of irradiation and poly(I:C) activation was studied in lung carcinoma cell lines, where, together with 4 Gy irradiation, it was demonstrated to enhance the cytotoxic effects of the monotherapies in a caspase-dependent manner *in vitro (39)* but no *in vivo* data were provided. Another study has demonstrated synergistic inhibition of tumor growth and enhanced induction of long-term immune memory cells in murine mammary and pancreatic carcinoma models using a combination of poly(I:C) injection with transplantation of alpha-emitting radiation seeds into the tumor (40), an experimental treatment that is currently being tested in clinical trials. However, unlike alpha-emitting radiation seeds, irradiation with a clinical linear accelerator, as used in our study, is a well-established treatment method for cancer patients.

Another interesting aspect of irradiation and immunity is that localized irradiation by itself, independent of additional innate immune activation, has been shown to improve tumor infiltration of adoptively transferred T cells in a pancreatic cancer model (42). With regard to irradiation intensity, other studies have shown that low doses (2–8 Gy) of irradiation elicit stronger antitumor immunity compared to higher doses, especially when given repetitively or when combined with other antitumoral treatments (13,43,44). In our study, despite the modest antitumoral response induced by 2 Gy irradiation alone, this low dose turned out to be more advantageous at co-activating RIG-I-mediated immunity than the higher doses (5 or 10 Gy).

Monotherapy with RIG-I agonists has been reported in several studies, demonstrating that intratumoral injection of RIG-I ligands induces an effective anti-tumor immune response (23,45). Importantly, our results highlight that RIG-I activation not only has the potential to improve the efficacy of conventional radiotherapy, but, at the same time, that RIG-I therapy itself can be improved by adding low-dose irradiation. According to our data, the combination with low-dose irradiation may enable a reduction in the required dose of RIG-I agonist to achieve effective treatment. To date, RIG-I agonist monotherapy has remained technically challenging and is limited by the injection volumes and RNA concentrations that can be achieved through the current delivery systems (46). Thus, if low-dose combination therapy reduced the amount of RIG-I ligand required, it could improve the feasibility of RIG-I agonist treatment.

Altogether, our study clearly demonstrates that the combination of DNA-damaging radiotherapy with innate-immune stimulating RIG-I ligand synergistically boosts p53-dependent immunogenic tumor-cell death, underscoring the rationale for evaluating a localized combination therapy that turns cold into hot tumors as an *in situ* cancer vaccine (27). Since melanoma is classically considered a “radioresistant” tumor, our study also provides a new rationale for reevaluating radiotherapy in combination with RIG-I activation for a broad range of cancers. Moreover, as with other synergistic combination treatments, it could potentially allow for a reduction of the individual radiation doses and thus reduce the severe side effects associated with standard radiotherapy.

## Authors’ contributions

Silke Lambing: formal analysis, investigation, writing –original draft, writing –review & editing, visualization

Stefan Holdenrieder: conceptualization, methodology, resources

Patrick Müller: investigation

Christian Hagen: investigation

Stephan Garbe: methodology, resources, writing review & editing

Martin Schlee: methodology, resources

Jasper G. van den Boorn: investigation, methodology, supervision, project administration

Eva Bartok: formal analysis, methodology, supervision, project administration, writing – original draft, writing –review & editing, visualization

Gunther Hartmann: conceptualization, funding acquisition, methodology, project administration, resources, supervision, visualization, writing –original draft, writing –review & editing

Marcel Renn: formal analysis, investigation, methodology, project administration, supervision, writing –original draft, writing –review & editing, visualization

## Acknowledgements

We thank Meghan Lucas for her critical reading of this manuscript. We thank Daniel Hinze for providing us with CRISPR gRNA/Cas9 plasmids targeting p53. We thank Jennifer Landsberg for her helpful scientific discussions.

---

**Supplementary Figure 1:**
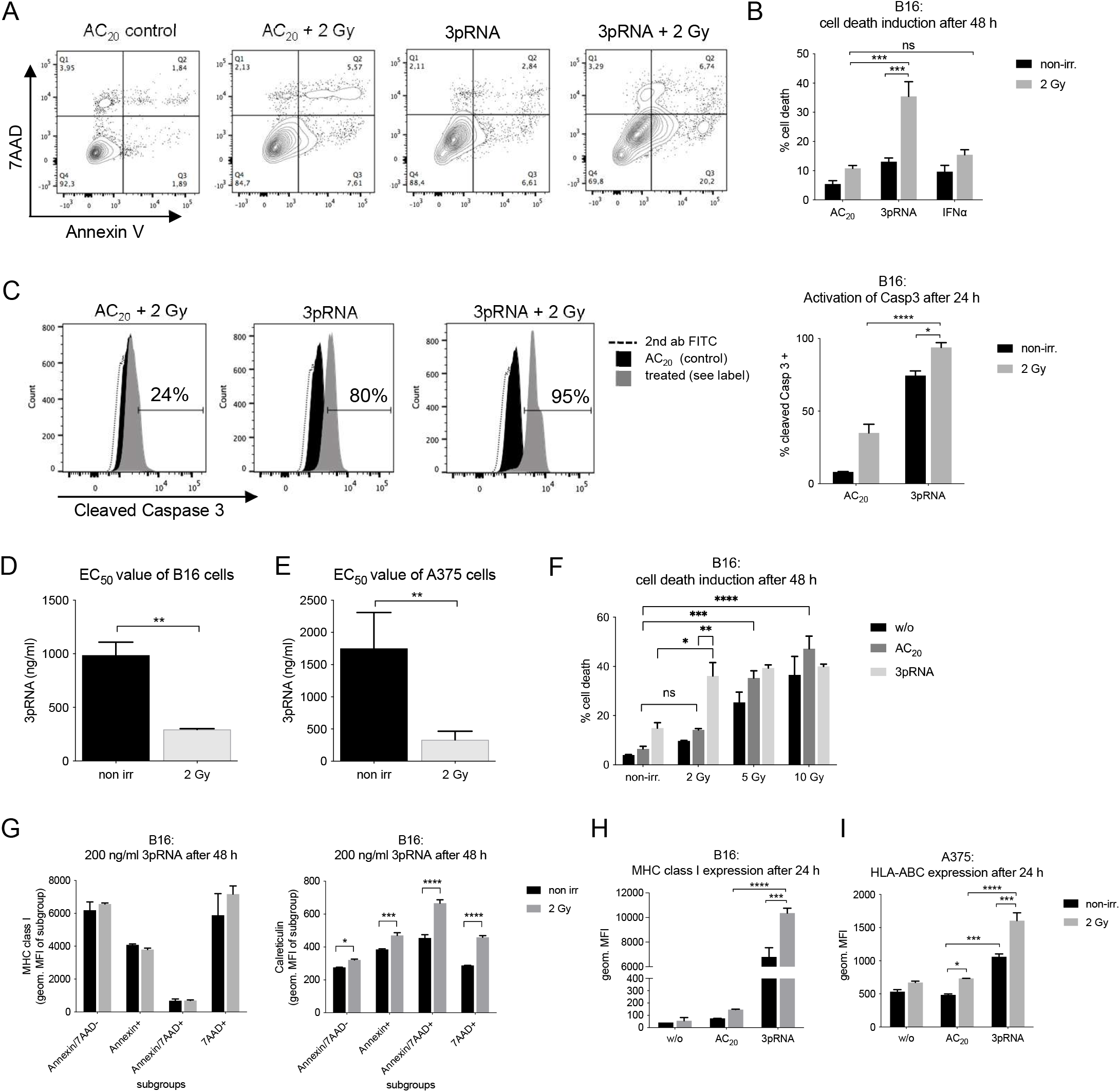
Irradiation enhances 3pRNA-induced immunogenic cell death in melanoma cells, as well as uptake by and co-stimulation of dendritic cells. B16 cells were transfected with 50 ng/ml 3pRNA or AC_20_ control RNA and simultaneously irradiated with 0 or 2 Gy. (A) Gating strategy Annexin V/7AAD staining. (B) Cells were additionally stimulated with 1000 U/ml recombinant IFNα and after 48 h, cell death was detected by Annexin V/7AAD staining. (C) Intracellular staining of activated, cleaved caspase 3 by fluorescently labeled antibody was measured after 24 h by flow cytometry. (D,E) Quantification of apoptosis induction by Annexin V/7AAD staining in B16 (D) and A375 (E) cells, 48 h after titration of 3pRNA concentration with and without 2 Gy, as shown in Fig. 1C,D. (F) B16 cells were transfected with 50 ng/ml 3pRNA and given different irradiation doses. 48 h later, cells were stained with Annexin V/7AAD and analyzed by flow cytometry. (G) Annexin V/7AAD staining after 48 h of 3pRNA (200 ng/ml) transfection and 2 Gy irradiation of B16 cells was combined with MHC class I and calreticulin fluorescent labeling. (H, I) MHC I expression on the surface of B16 (H) and A375 (I) cells 24 h after treatment, as detected by flow cytometry. (B−F) Mean and SEM, n=3. (G−I) Representative results with the mean and SD of n=3. * p<0,05; **p<0,01; ***p<0,001; ****p<0.0001; B, C, F, G, H; two-way ANOVA, D, E t-test. w/o: untreated, AC_20_: control RNA, 3pRNA: 5’-triphosphate RNA, non-irr: non-irradiated.

**Supplementary Figure 2:**
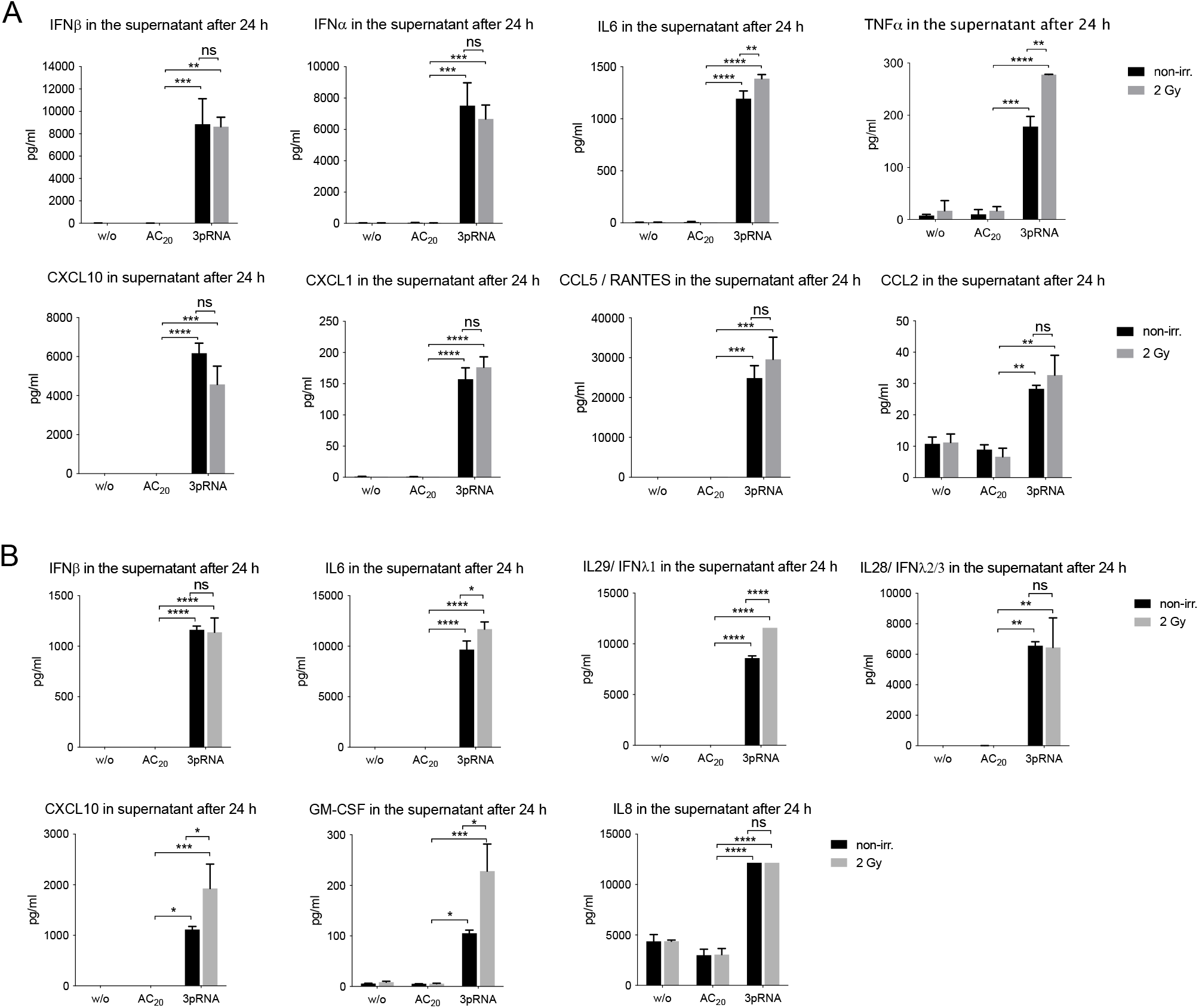
2 Gy irradiation has only minor influence on 3pRNA-induced cytokine release. Melanoma cells were transfected with 50 ng/ml 3pRNA or AC_20_ control RNA and simultaneously irradiated with 0 or 2 Gy. Supernatants were collected 24 h after treatment of B16 (A) and A375 (B) cells, and were analyzed by flow-cytometric multiplex analysis to detect different cytokines and chemokines. Shown is the mean and SD of one experiment with biological replicates measured in technical replicates. Not detected: B16 (A): IL10, GM-CSF, IL1b, IFNg, IL12p70; A375 (B): TNFa, IFNa2, IL10, IL1b, IFNg, IL12p70. * p<0,05; **p<0,01; ***p<0,001; ****p<0.0001; two-way ANOVA. w/o: untreated, AC_20_: control RNA, 3pRNA: 5’-triphosphate RNA, non-irr: non-irradiated.

**Supplementary Figure 3:**
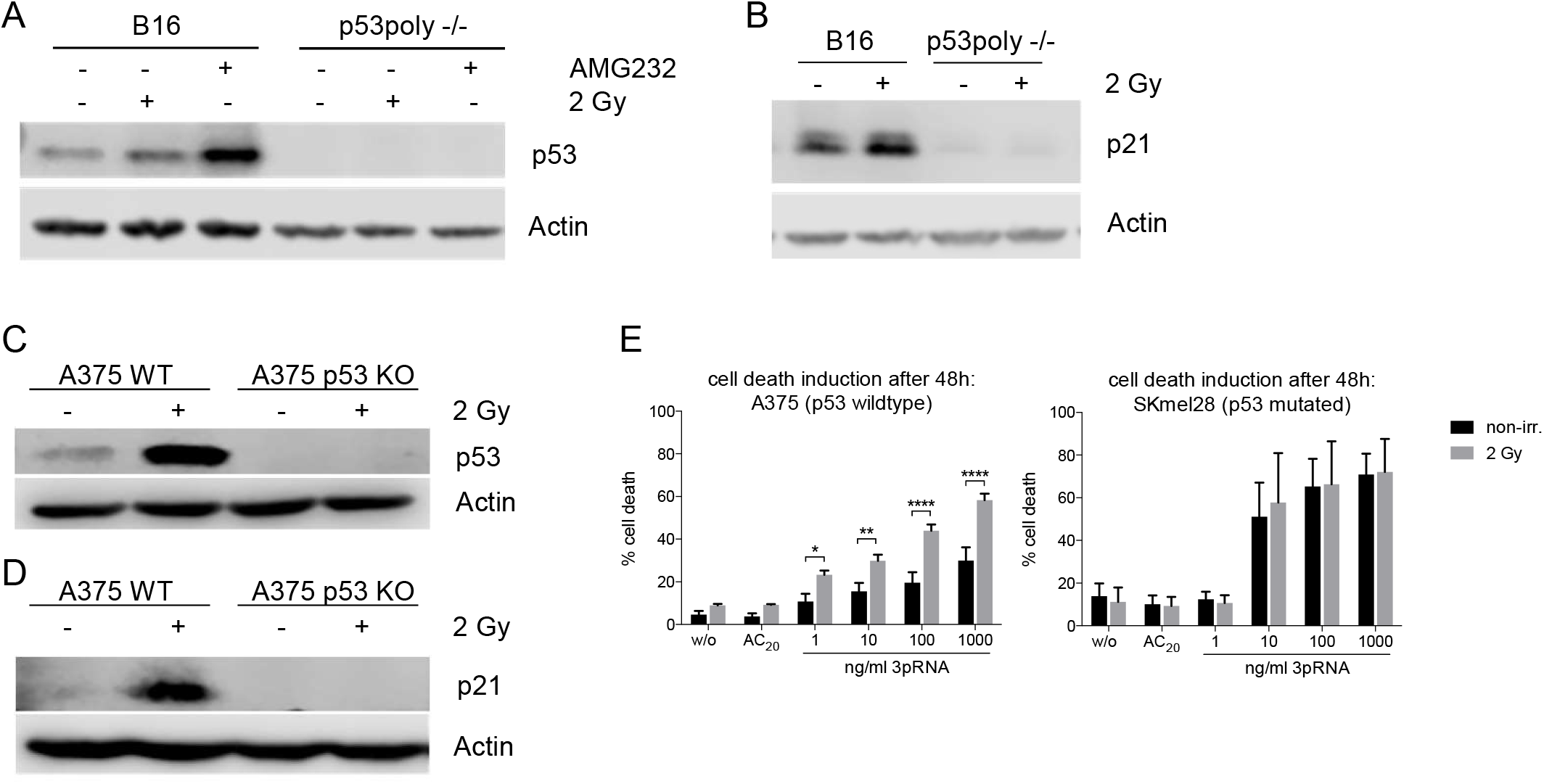
Establishment of p53 polyclonal knockout melanoma cells and functional comparison. Immunoblot analysis of p53 2 h (A, C) and p21 24 h (B, D) after irradiation with 2 Gy or treatment with 10 μM AMG232 in B16 and A375 wildtype and p53 polyclonal KO cells as indicated. Actin served as a loading control. (E) Human melanoma cell lines A375 and SKmel28 were transfected with increasing concentrations of 3pRNA and additionally irradiated with 2 Gy. Cell death was quantified 48 h later using Annexin V/7AAD staining and flow cytometry. Mean and SEM are shown from 3 independent experiments. p53 polyclonal knockout cells were generated by using the CRISPR/Cas9 system. * p<0.05; **p<0.01; ****p<0.0001; two-way ANOVA. AC_20_: control RNA, 3pRNA: 5’-triphosphate RNA, non-irr: non-irradiated

**Supplementary Figure 4:**
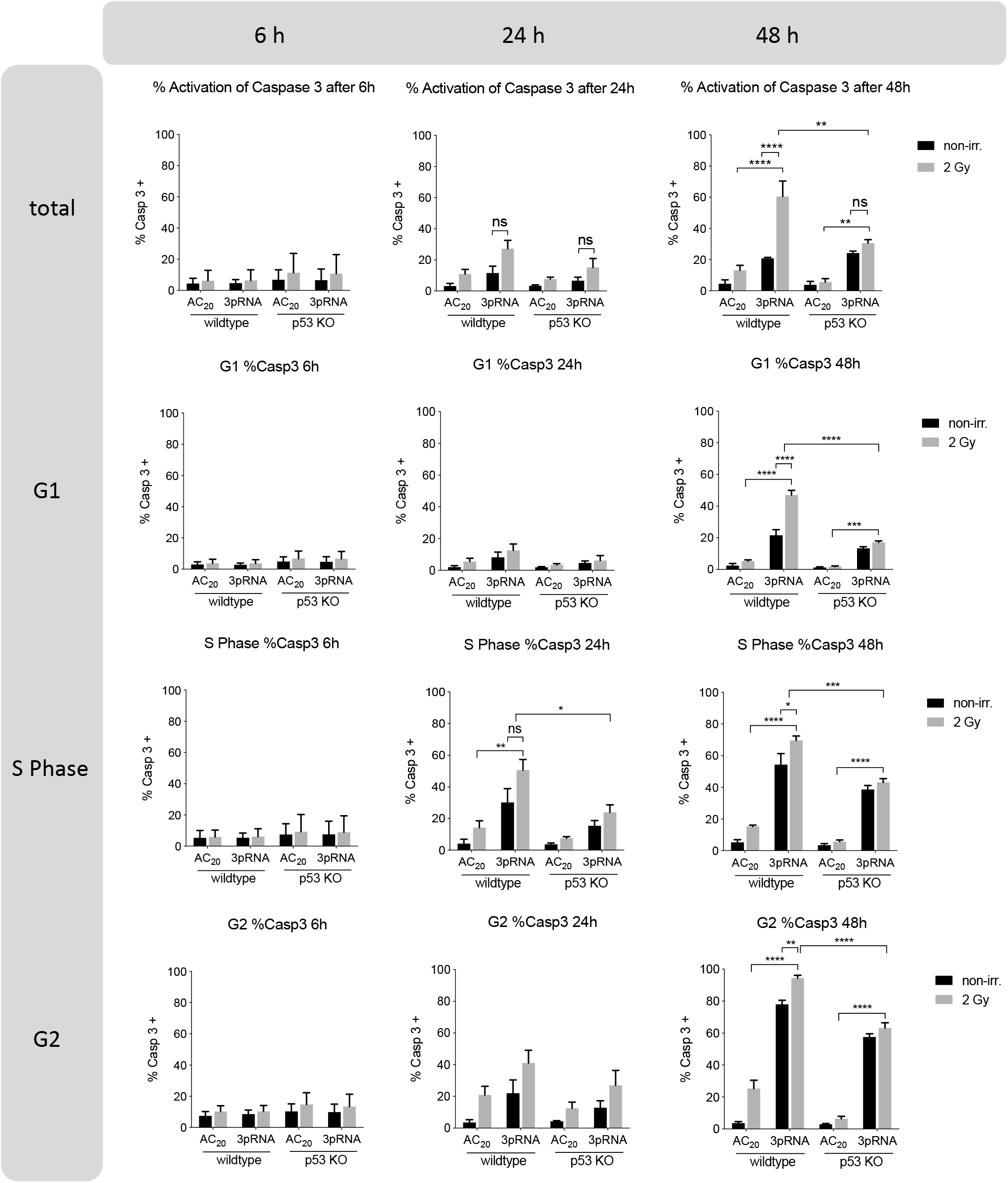
Increased cell death correlates with prolonged G2/M cell cycle in combinatorial RIG-I radio-immunotherapy. Flow-cytometric cell-cycle analysis of B16 cells treated with 50 ng/ml 3pRNA and 2 Gy irradiation using genomic Hoechst 33342 stain, in combination with intracellular staining with a caspase 3/7 cleavable dye at the indicated time points. * p<0,05; **p<0,01; ***p<0,001; ****p<0.0001; two-way ANOVA. ns: not significant, AC_20_: control RNA, 3pRNA: 5’-triphosphate RNA, non-irr: non-irradiated

